# *In vitro* selection of Engineered Transcriptional Repressors for targeted epigenetic silencing and initial evaluation of their specificity profile

**DOI:** 10.1101/2022.08.30.505942

**Authors:** Alessandro Migliara, Martino Alfredo Cappelluti, Sara Valsoni, Ivan Merelli, Davide Cittaro, Angelo Lombardo

## Abstract

Gene inactivation is instrumental to study gene function and represents a promising strategy for the treatment of a broad range of diseases. Among traditional technologies, RNA interference suffers from partial target abrogation and requirement for life-long treatments. On the other hand, artificial nucleases can impose stable gene inactivation through induction of DNA Double Strand Breaks, but recent studies are questioning the safety of this approach. Targeted epigenetic editing via Engineered Transcriptional Repressors (ETRs) may represent a solution, as a single administration of specific ETRs combinations can lead to durable silencing without DNA breaks induction. ETRs are proteins containing a programmable DNA Binding Domain (DBD) and effectors from naturally occurring transcriptional repressors. Specifically, a combination of three ETRs equipped with the KRAB domain of human ZNF10, the catalytic domain of human DNMT3A and human DNMT3Lwas shown to induce heritable repressive epigenetic states on the ETR-target gene. The hit-and-run nature of this platform, the lack of impact on the DNA sequence of the target, and the possibility to revert on-demand the repressive state by DNA demethylation, make epigenetic silencing a game-changing tool. A critical step is the identification of the proper position on the target gene where to tether the ETRs in order to maximize on target and minimizing off-target silencing. Performing this step in the final *ex vivo* or *in vivo* preclinical setting can be cumbersome. Taking CRISPR-Cas9 system as a paradigmatic DBD for ETRs, here we describe a protocol consisting in the *in vitro* screen of gRNAs coupled to the triple ETRs combination for efficient on-target silencing, followed by evaluation of the genome-wide specificity profile of top hits. This allows to reduce the initial repertoire of candidate gRNAs to a short list of promising ones, whose complexity is suitable for their final evaluation in the therapeutically relevant setting of interest.

## INTRODUCTION

Gene inactivation has traditionally played a key role to study gene function in both cellular and animal models. Furthermore, in the last two decades, with the raise of gene therapy, it has been proposed as a potentially game-changing approach to treat diseases caused by gain-of-function mutations^1^, infectious diseases^2^ or pathologies in which silencing of one gene may compensate for an inherited defect in another one^3^. Finally, genetic inactivation of key regulators of cell fitness and functional control has been proposed to enhance the efficiency of cell products for cancer immunotherapy^4^ and regenerative medicine^5^.

Among the different technologies to accomplish gene inactivation, one of the most promising is targeted epigenome editing^6,7^. At the core of this technology are the so called Engineered Transcriptional Repressors (ETRs), chimeric proteins consisting of a programmable DNA Binding Domain (DBD) and an Effector Domain (ED) with epigenetic repressive function. Zinc Finger Proteins (ZFPs)^8^-, Transcription Activator-like Effectors (TALEs)^9^ or CRISPR/dCas9^10^-based DBDs can be designed to selectively tether the ED to the promoter/enhancer sequence of the target gene to be silenced. Once there, the ED of the ETR performs its silencing activity by imposing heterochromatin-inducing repressive epigenetic marks such as histone modifications (H3K9^11, 12^ or H3K27^13^ methylation, H3 or H4 deacetylation^14^) and CpG DNA methylation^15^, according to the repressive domain used. In particular, inspired by the molecular processes of permanent transcriptional repression of endogenous retroviruses occurring in the pre-implantation embryo^16^, we and others have generated a combination of three ETRs exploiting the following EDs: i) the Krüppel-associated box (KRAB) domain of human ZNF10; ii) the catalytic domain of human *de novo* DNA methyltransferase 3A (DNMT3A); iii) the full-length human DNA methyltransferase 3-like (DNMT3L). KRAB is a conserved repressive domain shared by several Zinc Finger Proteins in higher vertebrates^17,18^ whose silencing activity is mainly based on its ability to recruit KAP1^19^, a scaffold protein that then interacts with several other heterochromatin inducers^20^, comprising the nucleosome remodeling and deacetylation (NuRD) complex^21^, the H3K9 histone methyltransferase SETDB1^22^, and the H3K9 methylation reader HP1^23,24^. On the other hand, DNMT3A actively transfers methyl groups on the DNA at CpG sequences^25^. The catalytic activity of DNMT3A is enhanced by its physical association with DNMT3L, an embryo- and germ cells-restricted paralogue of DNMT3A lacking the catalytic domain responsible for methyl group transfer^26,27^. DNA methylation at CpG-rich region - referred to as CpG islands, or CGI for short - embedded in the promoter/enhancer elements of mammalian genes is usually associated to transcription silencing^28^. Importantly, once deposited, CpG methylation can be stably inherited throughout mitosis by a UHRF1-DNMT1-based molecular complex^29^.

Stable over-expression of the ETRs in the target cell can be problematic, likely because of the increasing risks of off-target activity and squelching of endogenous interactors from their physiological target sites overtime. On the other hand, transient expression of single ETRs moieties can fail to induce long-term silencing at high efficiency^30^, hampering their therapeutic application. Therefore, a seminal breakthrough in the field was the evidence that the combination of the three KRAB-, DNMT3A- and DNMT3L-based ETRs can synergize and, even when only transiently co-delivered, impose on the promoter sequence of the target gene H3K9 and CpG methylation that are then read and propagated by the cell throughout mitosis leading to heritable silencing in multiple human and murine cell lines^30^. Of note, the epigenetic silencing imposed by the ETRs can be reverted on demand by targeted or pharmacological DNA demethylation^30^, a potential antidote in case of ETRs-related adverse events. All-in-one ETRs bearing the three KRAB-, DNMT3A- and DNMT3L-based EDs were also described, showing significant silencing efficiencies in human primary cells and cell lines^31,32^ and, in the latter case, showing to be effective against the large majority of protein-coding genes, with a wide permissive targeting window across promoters.

Furthermore, several studies employing the ETRs reported a high safety profile, with no major off-target activity detected in terms of *de novo* CpG methylation or alteration of chromatin accessibility genome-wide^30,31,32^. However, a dedicated analysis of the specificity profile of ETRs equipped with newly designed DBD should be recommended before clinical applications.

From a clinical perspective, targeted epigenetic silencing may provide critical advantages to both RNA interference-based knockdown^33^ and artificial nucleases-based gene disruption^8^ : in contrast to RNAi, targeted epigenome editing may induce full abrogation of its target *per cell* and does not require periodic treatment to ensure long-term silencing; in contrast to gene disruption, it leaves unaltered the DNA sequence, avoiding the generation of DNA double strand breaks that can then induce apoptosis and cell cycle arrest, potentially leading to a selection against cells with a functional p53 pathway^34,35^ and, especially in multiplex gene editing settings, chromosomal rearrangements^35^. Furthermore, by relaying on the irreversible mosaic outcome of Non-Homologous-End-Joining-mediated DNA double strand breaks repair^36^, gene disruption cannot avoid in-frame repair of the target into functional coding sequences as one of the final outcomes and, in contrast to epigenetic silencing, cannot be erased on demand. Finally, epigenome editing holds the potential to broaden the range of targetable genetic elements to classes fully or at least partially refractory to RNAi and gene disruption, such as non-transcribed regulatory elements and non-coding RNAs^30,32^.

The first critical step for any targeted epigenetic silencing application is to design a panel of ETRs covering the different regulatory sequences of the target gene and identify the best performing ones. The number of ETRs to be tested can be potentially significant, considering the increasing portion of the genome that can be targeted by the programmable DNA binding technologies constantly under development^37^. Performing the screen of the ETRs directly on the cell type in which to therapeutically silence the target gene would represent the most reliable option. However, high throughput screens can be technically cumbersome in primary cells due to their limited survival in culture and their often-suboptimal engineering capacity. Large-scale screens can be even more unfeasible *in vivo*. A more practical alternative consists in performing an initial screening of a large panel of ETRs in easily engineerable cell lines at first and then only validate the most promising ones in the therapeutically relevant cell type. A parallel issue is the selection of an appropriate read-out to measure the silencing efficiency of the ETRs. Directly assessing the transcript or protein levels of the target gene by RT-qPCR, Western blot or ELISA can be costly and time-consuming and may lack sufficient sensibility, thus limiting their application at high throughput scales. The generation of *ad hoc* engineered reporter cell lines in which a fluorophore is placed under the transcriptional control of the regulatory sequences of the target gene allows exploitation of flow cytometry-based approach to read epi-silencing at the single-cell level and at high-throughput pace.

Following these general considerations, here we will describe a protocol consisting in the *in vitro* arrayed screen of ETRs for on-target silencing efficiency followed by evaluation of the genome-wide off-target activity of the top hits. This workflow allows to shrink the initial repertoire of candidate ETRs to a short list of promising ones whose complexity is suitable for their final evaluation in the therapeutically relevant cell type of interest.

## PROTOCOL – INDEX

1. Generate a fluorescence-based reporter cell line to monitor the transcriptional activity of your target gene of interest by flow cytometry
2. Design gRNAs against your target gene for CRISPR/dCas9-based epigenetic silencing
3. Transient delivery of CRISPR/dCas9-based ETRs in the reporter cell line
4. Analyze the transcriptional activity of your target gene overtime to assess epi-silencing efficiency and durability
5. Evaluate specificity of the ETR treatment by and RNA-Seq and MeDIP-Seq

### 1. Engineer a fluorescence-based reporter cell line to follow the transcriptional activity of your target gene by flow cytometry

1.1 Identify cell lines expressing your target gene of interest. Browse the gene that you want to silence in the Human Protein Atlas^38^ and navigate through its “Cell line” section to identify those representative of your somatic target tissue (e.g. a hepatic cell line if your final targets are liver hepatocytes). Alternatively, it is possible to interrogate publicly available RNAseq database (i.e. NCBI GEO).
1.2 Among the candidates, prioritize cell lines for which efficient transient delivery protocols are available. Among the different modalities, the nucleofection represent the best option as it ensures high transfection efficiencies. On this regard, *Lonza* provides a database of the cell types for which plasmid DNA or mRNA nucleofection protocols have been optimized. Here we choose human erythroleukemia K-562 cells to generate a cell line reporting for the transcriptional activity of the *B2M* gene
1.3 Clone a donor template for the homologous recombination-mediated integration of a fluorescent reporter under the transcriptional control of your target gene (*Figure 1*).
  ⮚ Identify the target splicing isoform preferentially used in your selected cell line
  ⮚ Identify the region of your target gene where to integrate the fluorophore expression cassette
    ⮚ Recommended: either inside the first intron of the target gene (as in the example here shown: targeting tdTomato in the first intron of the human *B2M* gene) or just upstream of its stop codon.
    ⮚ Avoid targeting transcriptionally relevant elements such as CpG islands and regions enriched for H3K27 acetylation (marker of active promoters and enhancers).
  ⮚ Identify a gRNA that can cut effectively and specifically inside your target region. *Chopchop*^*39*^ is a valid and user-friendly online gRNAs selection tool
  ⮚ Design a donor template for your gRNA cut site. Donor templates consist of: a) a left homology arm (*n* base pairs (bp) matching the region just upstream of your gRNA cut site); b) a promoter-free transgene expression cassette (in the case here shown, a Splice acceptor site – 3X Stop Codon – IRES – tdTomato – BGH poly(A) favoring splicing with the first intron of the human *B2M* gene); c) a right homology arm (*n* bp matching the region downstream of your gRNA cut site). The length of the homology arms necessary to effectively induce homologous recombination can vary between different cell types (100-500 bp is a proper range for K-562 cells).
1.4 Deliver CRISPR/Cas9 and the donor template inside your target cell line. Choose a delivery system representing a good compromise between transgene delivery efficiency, toxicity and costs. For K-562 Lonza-mediated nucleofection of plasmids separately encoding for the donor template, Cas9 and the gRNA represents a valuable strategy since plasmids can be easily produced and delivered at high copies in these cells.
1.5 Monitor the expression of the fluorescent reporter in treated cells overtime by flow cytometry
1.6 Wait at least two weeks to allow dilution of non-integrated copies of the donor template. Despite standard promoter-less, the donor template – especially if plasmid-based – may contain cryptic promoter sequences inside it, leading to transgene expression from non-integrated donor copies
1.7 Clone reporter-positive cells through Fluorescence Activated Cell Sorting (FACS) at a single cell level
1.8 Screen reporter-positive clones by PCR to select one bearing a bi-allelic integration of the reporter inside your target locus

**Figure 1.**
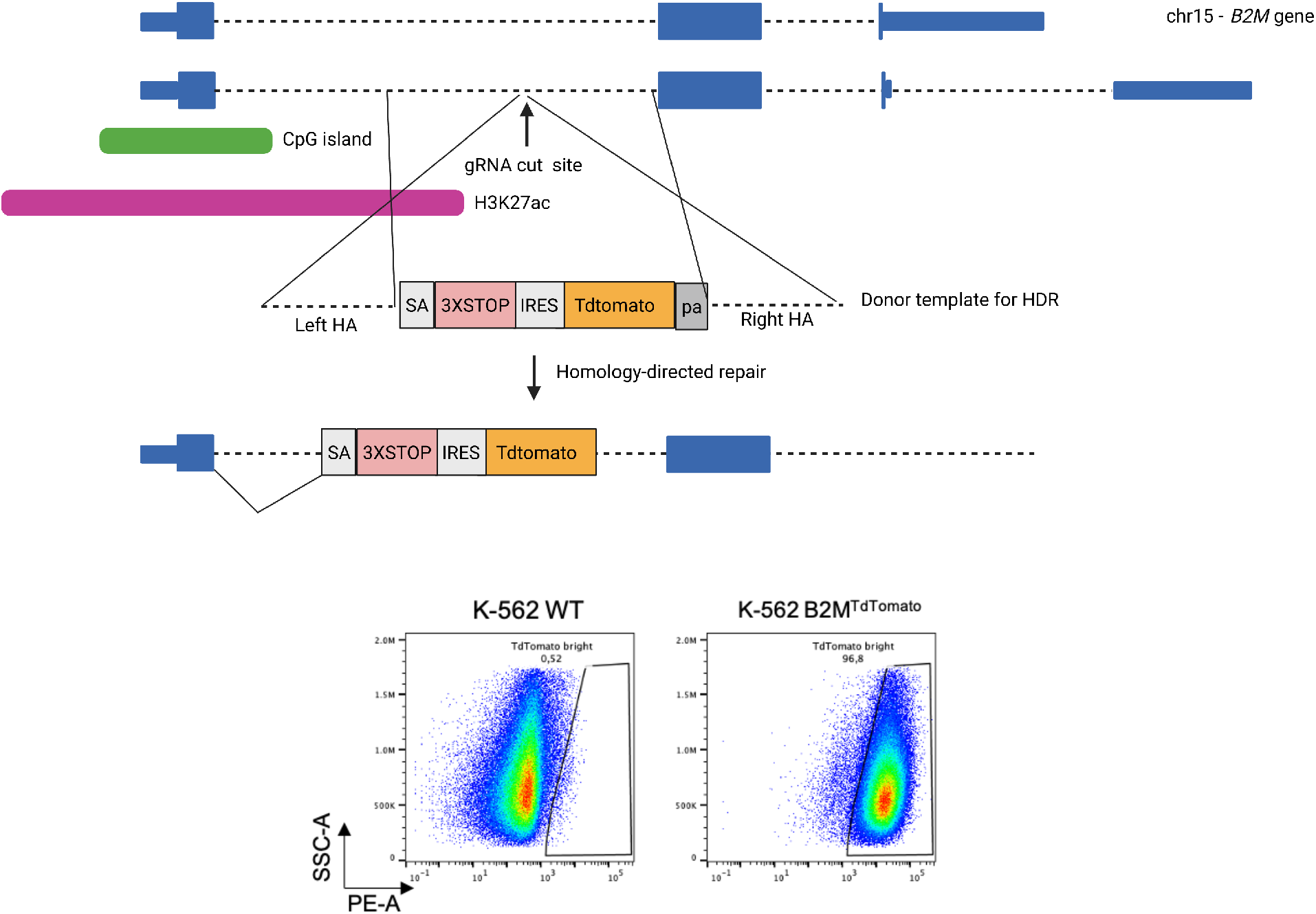
Integration of a fluorescent reporter under the regulatory elements of the target gene by homologous recombination. Top: Schematics of strategy to integrate a Tdtomato fluorescent reporter in the first intron of the B2M gene by CRISPR/Cas9-induced Homology-directed repair. Bottom: representative dot plots of K-562 cells pre- and post-integration of the TdTomato reporter in the first intron of the B2M gene. Bottom: K-562 cells pre- and post- Homologous recombination-mediated integration of TdTomato inside the first intron of the B2M gene.

### 2. Design gRNAs for CRISPR/dCas9-based epigenetic silencing of your target gene

2.1 Browse your gene in the *UCSC genome browser*^40^ and extract the nucleotide sequence of regions potentially regulating its transcriptional activity, such as CpG islands and sites enriched for H3K27 acetylation (marker of active promoters and enhancers).
  ⮚ According to a recent study, the best targeting region is a 1kb window centered on the Transcription Start Site of the target gene^32^
2.2 Paste the selected sequences in the *Chopchop* online tool and select “repression” as purpose of the gRNAs to be retrieved. *Chopchop* will then provide a list of gRNAs mapped on the genetic sequence of interest and listed according to a score considering both the number of off-target matches and the predicted on-target efficiency (Fig.2).
2.3 Select at least 20 gRNAs per each of your target sequences. If possible, try to select equally spaced gRNAs with no full matches with other intragenic sequences throughout the genome.

**Figure 2.**
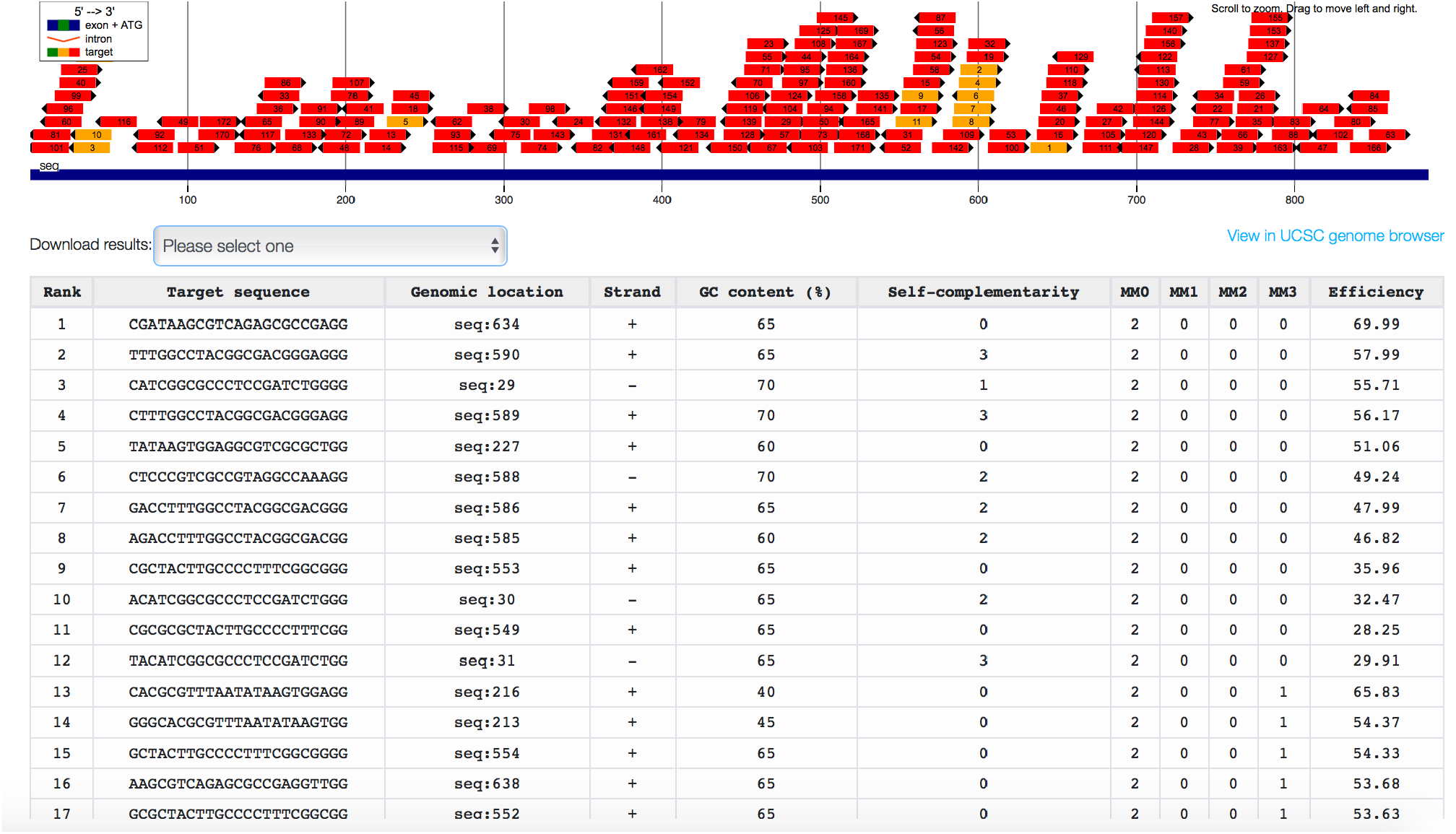
*In silico* identification of gRNAs matching your target sequence, ranked for predicted efficiency and specificity. Chopchop’s output interface showing gRNAs targeting the CpG Island embedded in promoter sequence of the B2M gene, ranked for both number of off-target sequences with 0 (MM0), 1 (MM1), 2 (MM2) or 3 (MM3) mismatches and the predicted on-target efficiency.

### 3. Arrayed transient delivery of CRISPR/dCas9-based ETRs in the reporter cell line

3.1 Among the transgene delivery systems previously reported for your target cell line, choose one allowing only transient transgene expression, maximizing delivery efficiency and minimizing cell manipulation-related toxicity.
  ⮚ In the case of K-562 cell, both plasmid and mRNA nucleofection are highly recommended, with plasmid production representing an easier and cheaper alternative.
3.2 Clone both the gRNAs selected in step 2 and CRISPR/dCas9-based ETRs in the transgene delivery system of choice. In case of plasmid-mediated delivery:
  ⮚ A plasmid encoding for an all-in-one ETR, termed CRISPRoff-v2.1^32^ is available on Addgene (Addgene No. 167981)
  ⮚ Plasmids separately encoding for dCas9:KRAB, dCas9:DNMT3A and dCas9:DNMT3L^30^ are not available on Addgene. They were cloned by replacing the VP160 trans-activator from the plasmid pAC154-dual-dCas9VP160-sgExpression^41^ (Addgene No. 48240) with either the KRAB, DNMT3A or DNMT3L domain-coding sequence^30^.
  ⮚ Transform plasmids encoding for the ETRs in One Shot™ TOP10 Chemically Competent E. coli cells (for the transformation procedure, follow the instructions of the TOP10 vendor). Screen the colonies for the presence of the ETR-bearing plasmid by either restriction enzymes digestion or Sanger sequencing and finally choose one of the positive colonies for plasmid DNA Midiprep production (follow the instructions of the vendor).
  ⮚ A plasmid for the human U6 promoter-driven expression of gRNAs (phU6-gRNA^42^) is available on Addgene (Addgene No. 53188). In order to clone the gRNAs you selected inside this backbone follow this protocol (Fig.3):
    3.2.1 Append a 5’-CACCG-3’ sequence upstream of the protospacer (the first variable 20-nt of your gRNA) → generate a 25-nt SGfw oligo
    3.2.1 Append a 5’-AAAC-3’ sequence upstream of the reverse complement of the protospacer and a 5’-C-3’ downstream of it → generate a 25-nt SGrv oligo
    3.2.2 Order both the SGfw and SGrv oligos as salt-free, water resuspended at a 100μM concentration
    3.2.3 Add 1 ul of each oligo to 2 ul of annealing buffer (10mM Tris, pH7.5-8.0, 50mM NaCL, 1mM EDTA) and 16 ul of water
    3.2.4 Perform oligos annealing by placing the solution in a thermocycler programmed to start at 95°C for 10 minutes. Then, gradually cool to 25°C over 45 minutes.
    3.2.5 Dilute 1μL of annealed oligos with 99μL nuclease-free water and then ligate 1 ul of this dilution with 50 ng of phU6-gRNA plasmid previously digested with the BbsI restriction enzyme (follow the instructions of BbsI and Ligase vendors for the digestion and ligation procedure)
    3.2.6 Transform 20 μL One Shot™ TOP10 Chemically Competent E. coli cells with 2 μL of the ligation product (follow the instructions of the TOP10 vendor for the transformation procedure)
    3.2.7 Pick multiple colonies for the plasmid DNA Miniprep production (follow the instructions of the vendor) and control the successful cloning of the protospacer by Sanger sequencing with the following primer matching to the U6 promoter 5’-GAGGGCCTATTTCCCATGATT-3’.
    3.2.8 Choose one of the positive colonies for plasmid DNA Midiprep production (follow the instructions of the vendor).
  3.3 Deliver the selected gRNAs and CRISPR/dCas9-based ETRs in the reporter cell line transiently in array (Fig.3). Here, as representative case, the workflow for the nucleofection of plasmids encoding for dCas9:KRAB, dCas9:DNMT3A, dCas9:DNMT3L and gRNAs targeting the B2M CpG island in B2M^TdTomato^ K-562 cells is shown:
    3.3.2 Prepare separate tubes, each containing 500 ng of dCas9:KRAB-, dCas9:DNMT3A- and dCas9:DNMT3L-enconding plasmids but differing for the gRNA to be tested (125 ng of a gRNA-encoding plasmid per tube). Include a gRNA-free condition as negative control
    3.3.3 Pellet 5*10^5 B2M^TdTomato^ K-562 cells per tube and resuspend them in SF nucleofection solution supplemented with the Supplement 1 provided by Lonza.
    3.3.4 Mix the resuspended cells and the plasmids mix and perform nucleofection according to the K-562-optimized Amaxa™ 4D-Nucleofector™ Lonza program (FF-120)
    3.3.5 Resuspend the cells in 200 ul of previously warmed RPMI-1640 mammalian cell culture media and place them back in the incubator

**Figure 3.**
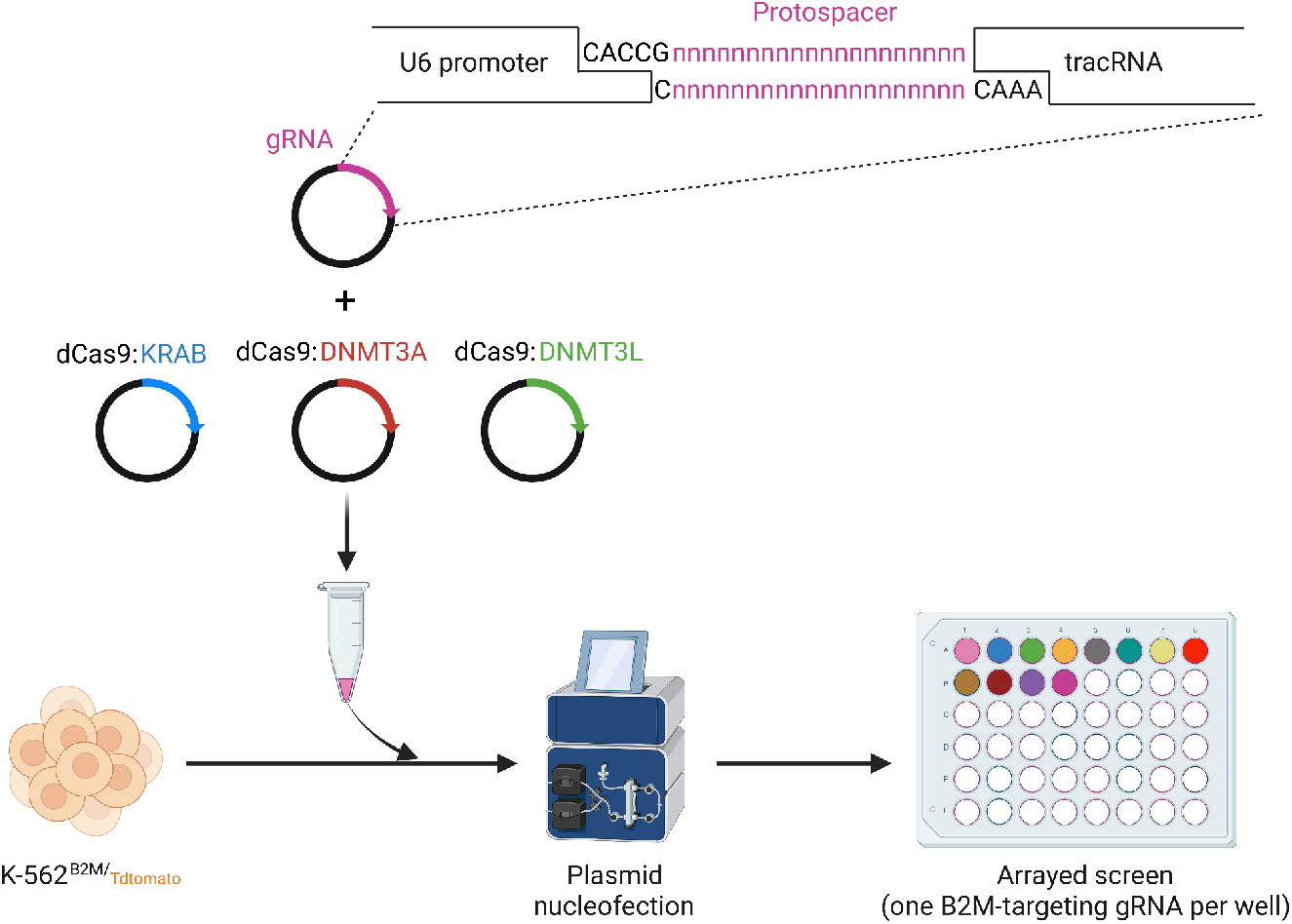
Cloning and arrayed nucleofection of gRNAs for dCas9 ETRs-mediated epigenetic silencing. Top: oligo-mediated protospacer cloning in a human U6-gRNA expressing plasmid. Bottom: arrayed screen of B2M-targeting gRNAs for CRISPR/dCas9-based epigenetic silencing by plasmid nucleofection in K-562^B2M/TdTomato^ cells.

### 4. Analyze the transcriptional activity of your target gene overtime

4.1 Measure by flow cytometry the percentage of silenced cells at different time points after the delivery of the ETRs.
  ⮚ Use WT cells - not bearing the fluorophore coding sequence - to set the threshold of reporter-negative cells
  ⮚ Use edited cells transfected with ETRs and no gRNA to set the gate for reporter positive cells
  ⮚ Used edited cells transfected with Cas9 nuclease and a gRNA targeting the coding sequence of the fluorescent reporter as a genetic disruption control. This can be useful to both monitor the CRISPR delivery efficiency of the system and the fitness of target KO cells overtime: genetic disruption is by definition a stable event. Counterselection of target disrupted cells overtime would indicate that the target gene is essential for the physiology of the cell
  ⮚ Include both short- (day 3, day 7, day 10) and long-term (day 21, day 35) time points.
4.2 Identify the top 3 gRNAs in terms of long-term silencing efficiency. Select by Fluorescence-Activated Cell Sorting (FACS) the reporter-negative subpopulation stably maintained in those conditions for further analyses

### 5. Evaluate specificity of the ETR treatment by RNA-Seq and MeDIP-Seq

5.1 Evaluate genome wide transcriptional deregulation upon ETRs delivery by RNA-Seq
  5.1.1 For both the reporter-silenced subpopulation of the samples treated with the top 3 gRNAs and mock-treated cells extract RNA using the RNeasy Mini kit (QIAGEN) according to manufacturer’s instruction.
  5.1.2 Amplify cDNA from total RNA (using the Ovation Human FFPE RNA-seq Library System (Nugen)
  5.1.3 Fragment cDNA with E220 COVARIS ultrasonicator (Covaris)
  5.1.4 Prepare sequencing libraries (using the Ovation Human FFPE RNA-seq Library System (Nugen)
  5.1.5 Perform library quantification and quality control on Bioanalyzer2100 (Agilent).
  5.1.6 Pool, denature, and dilute barcoded libraries to a 7 pM final concentration
  5.1.7 Perform cluster formation on board of HiSeq 2500 Rapid Mode flow cell (Illumina).
  5.1.8 Perform Sequencing By Synthesis (SBS) according to HiSeq PE protocol v2 (Illumina) on HiSeq 2500 system (Illumina) set to 200 cycles, yielding an average of 30M reads/sample.
  5.1.9 Align read tags to appropriate reference genome (e.g. hg38 using STAR (v 2.3.0 or higher)^43^), with default parameters. Count features using Rsubread package^44^, with appropriate gene model (e.g. gencode^45^). Perform alignment on tdTomato sequence and quantify it separately.
  5.1.10 Normalize feature counts, summarized at gene level, with the R/Bioconductor package edgeR using the Trimmed Median of Means approach^46^. Use a filter of at least one count per million (cpm) in at least 3 samples to discard low-expressed genes.
  5.1.11 Evaluate differential gene expression with Negative Binomial Generalized log-linear model implemented in the R/Bioconductor package edgeR (function glmFit)^47^. Set a threshold of 0.01 on adjusted p-values (Benjamini-Hochberg correction, BH) to retain differentially regulated genes. Calculate RPKM and TPM using R.
5.2 Evaluate the off-target CpG methylation activity of the ETRs by MeDIP-Seq
  5.2.1 For both the reporter-silenced subpopulation of the samples treated with the top 3 gRNAs and mock-treated cells extract genomic DNA with QIAamp DNA Mini Kit (QIAGEN)
  5.2.2 Sonicate 500 ng of genomic DNA with E220 COVARIS ultrasonicator (Covaris) and prepare sequencing libraries with the NextFlex Methylseq kit 1 (Bioo Scientific).
  5.2.3 After adaptor ligation step, pool the samples and immunoprecipitate them using a monoclonal antibody directed against 5-methylcytosine (MagMeDIP kit, Diagenode).
  5.2.4 Purify enriched and control libraries (not immumoprecipitated) using the IPure kit (Diagenode). Evaluate the enrichment efficiency by performing quantitative real-time PCR with internal controls provided by the kit (spiked-in DNA from A. thaliana).
  5.2.5 Amplify the libraries according to the NextFlex Methylseq kit 1 protocol and perform a quality control on the amplification product on Bioanalyzer2100.
  5.2.6 Perform Sequencing of the libraries according to HiSeq PE protocol v2 (Illumina) on HiSeq 2500 system (Illumina) set to 200 cycles, yielding an average of 30M reads/sample.
  5.2.7 Align the sequencing read tags to the appropriate reference genome (e.g. hg38) using bwa (v 0.7.5 or higher)^48^ and then identify peaks using MACS (v 2.0.10 or higher)^49^ allowing for broad peaks identifi- cation (–slocal = 0,–llocal = 500000). Unify peaks obtained in different samples using bedtools multiintersection tool^50^ with clustering option. Calculate read counts over the final peak list using bedtools, discarding duplicated reads.
  5.2.8 Identify differential methylation by adopting the generalized log-linear model implemented in edgeR (function glmFit) and normalizing using Conditional Quantile Normalization^51^ in order to model region-wise GC-content.
    ⮚ To retain differentially methylated regions, we recommend to use a threshold of 0.01 on BH adjusted p-values
    ⮚ Perform the analysis of repeated sequences as follows:
      5.7.1 Filter MeDIP results for nominal p value < 0.01 and absolute logFC > 1 so that two datasets are produced (up/down)
      5.7.2 Count the number of intersections of the regions in these datasets with all classes of repeats annotated in the RepeatMasker track available for the chosen reference genome and express the count as ratio over the number of regions for each dataset.
      5.7.3 Also extract the ratio of methylome that overlaps each class of repeats and perform a Chi-square test to detect any significant enrichment.
5.3 Evaluate if differentially transcribed or differentially methylated regions between ETRs- and mock-treated samples map to *in silico*-predicted off-target gRNA binding.
  5.3.1 As off-target gRNA binding prediction tool use CRISPR design suite^52^
  5.3.2 For every putative off-target region look at the closest Transcription Start Site (TSS) and the closest methylated region
    ⮚ Consider a true off-target effect a region associated either to a gene regulated with <u>FDR</u> < 0.01 and a distance to TSS smaller than 10Kb or a methylated region regulated with FDR < 0.01 and a distance lower than 1 Kb.
  5.3.3 Identify the number and features of the regions transcriptionally altered or over-methylated in samples treated with each of your top 3 gRNAs compared to mock-treated samples (Fig.5) in order to identify the most specific among your gRNAs.
    ⮚ We recommend to rank the potential impact of an off-target site on the physiology of your target cells in the following order:
      1. Intragenic, regulatory region – physiologically expressed gene
      2. Intragenic, exonic region - physiologically expressed gene
      3. Intragenic, intronic region - physiologically expressed gene
      4. Intragenic, regulatory region –not expressed gene
      5. Intragenic, exonic region - not expressed gene
      6. Intragenic, intronic region - not expressed gene
      7. Intergenic region

## REPRESENTATIVE RESULTS

Upon delivery of a donor template for homologous recombination-mediated integration of the fluorescent reporter in your target locus coupled with the CRISPR/Cas9 system – e.g., by plasmid nucleofection in the case of K-562 cells -, you should note the appearance of a reporter-positive cells in your treated sample (Fig.1, Bottom). If this does not occur, check back the accuracy of design and cloning of both your donor template and CRISPR/Cas9 reagents. If confirmed, try to optimize the doses of reagents and the delivery protocol itself. For instance, for cell lines in which nucleofection has not been optimized yet, *Lonza* provides a kit for testing a panel of different protocols to identify the best performing one.

Once obtained your reporter cell line, select promoter/enhancer sequences of your target gene and design a panel of matching gRNAs. Exploit *in silico* prediction tools such as Chopchop to both identify the gRNAs sequence and rank them in terms of predicted efficiency and specificity (Fig.2). If no gRNAs are retrieved, control the presence of the canonical Cas9 PAM (NGG) in your target sequence. If no PAM sequences are present, consider to either shift to ETRs based on alternative PAM-independent Cas9 variants (no ETRs published with these variants yet) or to switch to alternative DNA binding Domain platforms such as Zinc Finger^53^ and TALE^30^. On the other hand, if only gRNAs with low predicted efficacy/specificity are retrieved, consider expanding your target DNA sequence and search for additional gRNAs.

Upon transient delivery of the triple ETRs combination (or CRISPRoff-v2.1) together with single gRNAs, by assessing the expression of your fluorescent reporter through flow cytometry you should note an acute silencing induction peak which is then at least partially reabsorbed due to mitotic dilution of the ETRs-encoding plasmids overtime (Fig.4C). If the ETRs/gRNA combination effectively deposits CpG methylation on the target locus, permanent repression of the reporter should occur in a sizable fraction of treated cells (Fig.4C). Different gRNAs may show variable long-term silencing efficiency (Fig.4B, C). If none of the gRNAs tested show the ability to instruct long-term silencing, consider increasing the amount of gRNAs and ETRs delivered and to test pools of your top gRNAs in terms of acute silencing induction activity. If you still fail to obtain long-term silencing, consider designing and testing additional gRNAs or to switch to different DNA Binding Domain platforms.

**Figure 4.**
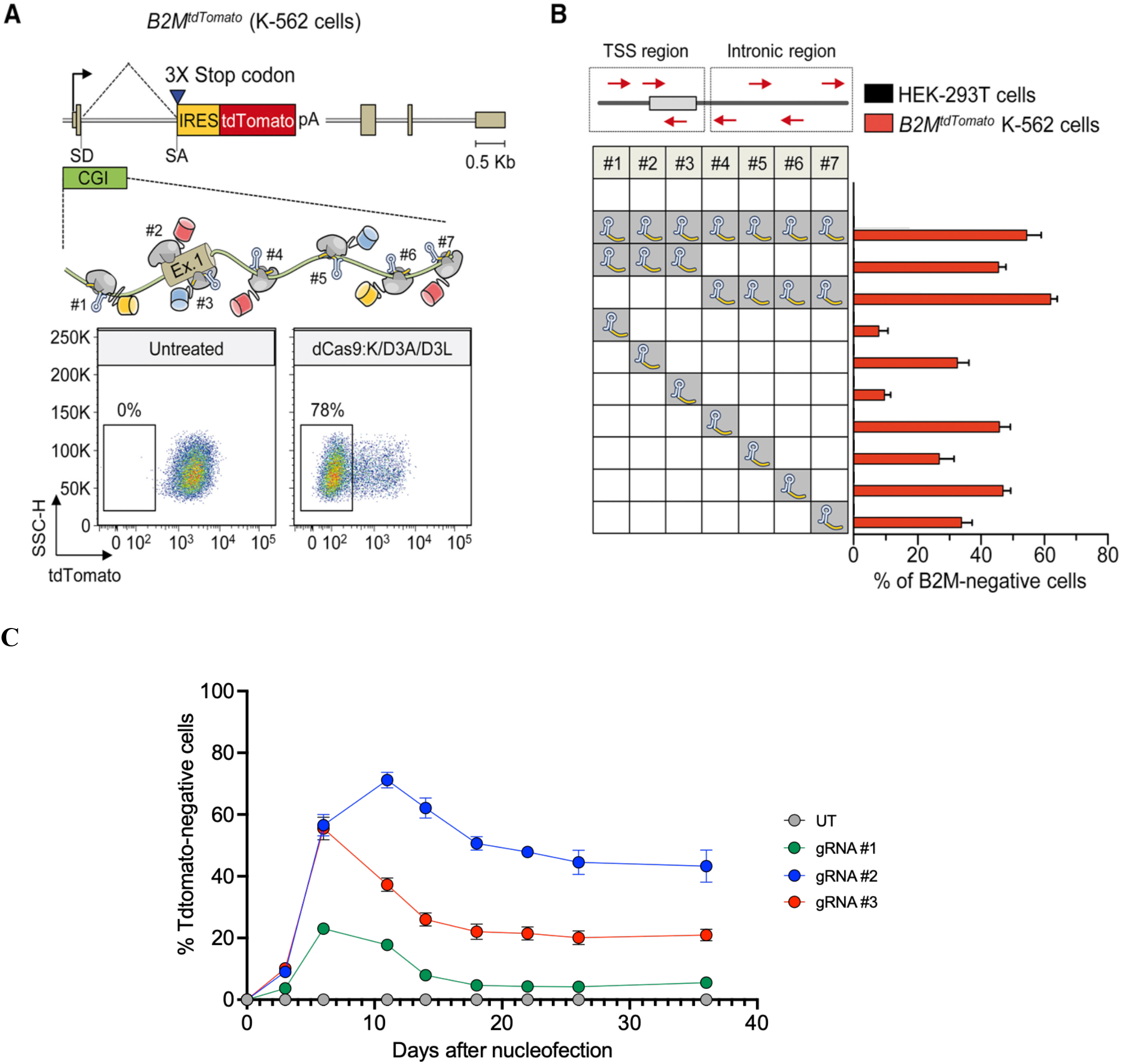
Screen for gRNAs effectively inducing long-term silencing of the target gene. A. Top: schematics of the B2M^tdTomato^ gene depicting in the enlarged area the relative order and orientation of binding of dCas9-based ETRs complexed with gRNAs. CGI, CpG island. Bottom: representative dot plots of B2M^tdTomato^ K-562 cells either before (left) or after (right) ETR silencing. Analyses at 30 days post-ETR transfection. B: silencing activity of the indicated gRNAs (either in pools or as individual gRNAs) targeting the CpG island of B2M (red arrows in the top schematic indicate orientation of the gRNAs) in K-562 B2M^tdTomato^ cells at day 30 post-nucleofection. Data show percentage of tdTomato negative cells (mean ± SEM; n = 4 independent transfections for each treatment condition; Adapted from Amabile, Cell, 2016). C: Time-course analysis of B2M^tdTomato^ K-562 cells upon transfection with plasmids expressing the triple ETRs combination and the indicated B2M CpG island-targeting gRNAs. Data show percentage of Tdtomato-negative cells (mean ± SEM; n = 3 independent transfections for each treatment condition).TSS, transcription start site. Unpublished data.

Identify by MeDIP-Seq the differentially methylated regions genome-wide between cells in which your target has been long-term silenced separately by your top 3 gRNAs and untreated cells.

Ideally, a highly specific gRNA should only induce a peak of *de novo* CpG methylation at your target site (Fig.5). If all of your top gRNAs show a low specificity profile, try if exposing the cells to lower amount of gRNAs and ETRs may reduce the off-target CpG methylation activity but still maintain acceptable on target silencing efficiency. If not, consider to characterize the off-target activity of the gRNAs ranked in a lower position in your on target efficiency list. If this fails too, consider to test ETRs delivery systems allowing shorter transgene persistence inside your cells (e.g. nucleofection of CRISPR/dCas9 Ribonucleoproteins instead of plasmids) and/or alternative DNA Binding Domain platforms such as Zinc Finger^53^ and TALE^30^.

**Figure 5.**
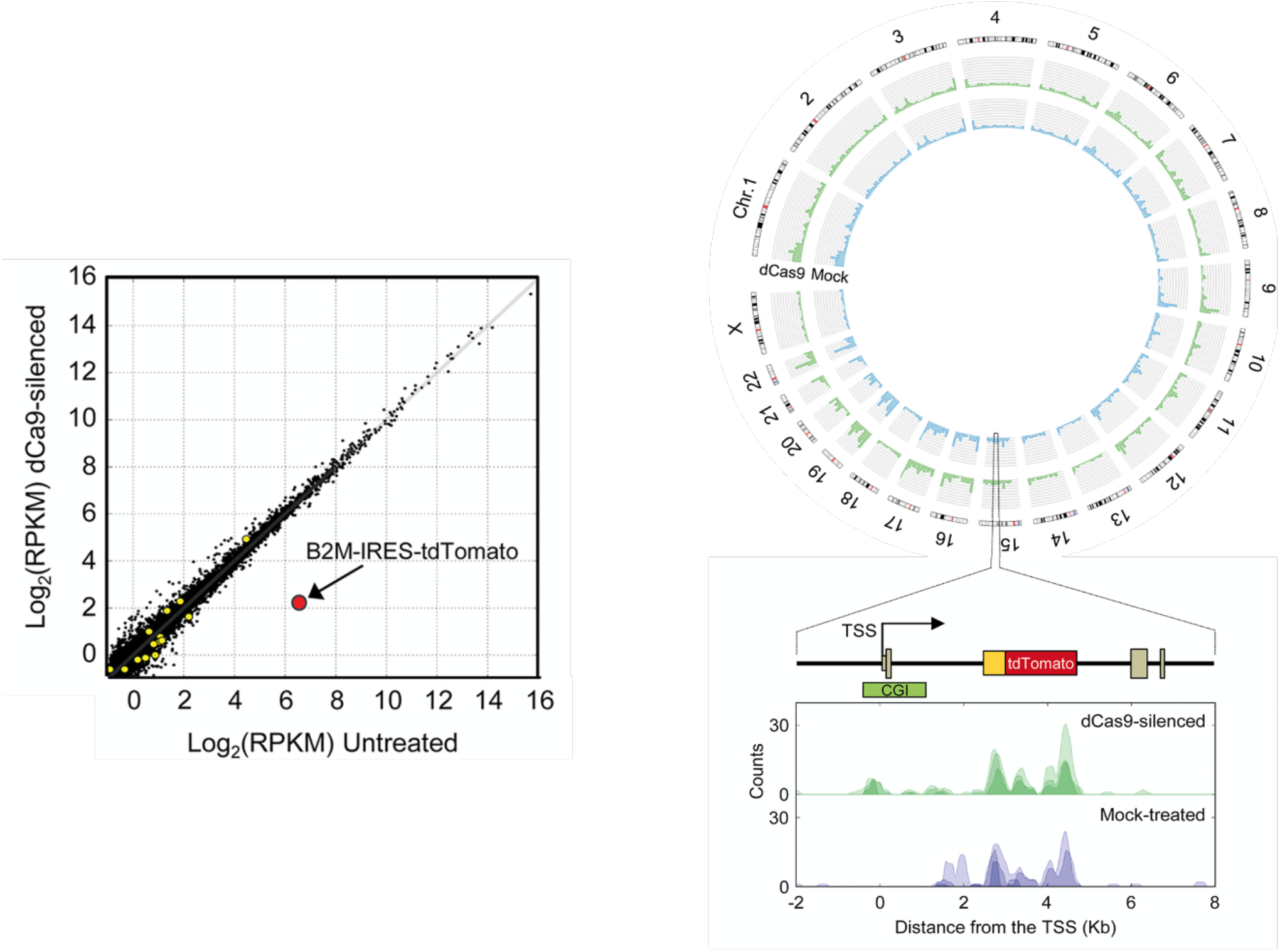
Genome-wide analysis of the ETRs specificity by RNA-Seq and by MeDIP-Seq. Left: Comparison of expression levels in mock-treated B2M^tdTomato^ K-562 cells and cells treated with the triple dCas9:KRAB, dCas9:DNMT3A, dCas9:DNMT3L ETRs combination and with a gRNA targeting the CpG island of the B2M gene. Values are expressed in log2 of read per kilobase per million (RPKM) of mapped reads. Black dots represent genes expressed at comparable levels in all conditions; yellow circles represent genes differentially regulated under a FDR < 0.01; red circle represents the B2M-IRES-tdTomato transcript. Top right: circos plot showing whole-genome MeDIP-seq profiles of mock-treated B2M^tdTomato^ K-562 cells (blue) or cells treated with the triple dCas9:KRAB, dCas9:DNMT3A, dCas9:DNMT3L ETRs combination and with a gRNA targeting the CpG island of the B2M gene (green). Bottom right: the methylation status of the B2M^tdTomato^ locus in the indicated samples is shown. Three replicates are represented in each pileup: pileup of aligned reads were smoothed using a Gaussian window. Adapted from Amabile, Cell, 2016

## DISCUSSION

Targeted epigenetic silencing may represent a promising solution to treat disorders that can benefit from permanent genetic inactivation, including diseases caused by gain-of-function mutations^1^, infectious diseases^2^ and pathologies in which silencing of one gene may either compensate for an inherited defect in another one^3^ or unleash the full potential of adoptive cell therapies^4,5^. One of the key, preliminary steps in any epigenetic silencing protocol is to identify the proper position on the target gene where to direct the ETRs that will deposit the repressive epigenetic marks necessary to turn off the transcriptional activity of the target.

Different transcriptional start site-proximal and -distal regulatory elements concur to support the transcriptional output of human genes^54^. And different target sites per each specific regulatory element can now be identified thanks to the growing number of programmable DNA binding technologies^6^. Therefore, protocols, as the one described here in engineered cell lines, can be used to nominate individual target sites and/or genomic regions amenable to ETR-mediated episilencing, before embarking in cumbersome and time-consuming evaluation efforts of best candidates in the final therapeutic setting.

Here we will further discuss some critical aspects of the protocol described above.

### Engineering of a reporter cell line predictive of the final therapeutic target

Despite the constant optimization of cell engineering protocols - here required to insert the cassette coding for the fluorescent reporter in the target gene and to deliver the ETRs -, it is not granted that they might be already available for the cell line that most resemble your final therapeutic target. In this case, different mitigation strategies can applied: a) take advantage of optimization kits provided by vendors to optimize *in house* the transfection protocol for your cell line of interest; b) switch to other cell lines still expressing your gene of interest, but belonging to tissues other than your final therapeutic target - for which engineering protocols have been extensively optimized. In case these options are not available, you might consider to either switch to primary cell types or organoids representing your final target.

### Evaluate the on-target silencing efficiency of the ETRs

Once a cell line reporting for the transcriptional activity of your target gene has been generated, you are ready to test the on-target, long-term silencing efficiency of ETRs equipped with your panel of *in silico* selected gRNAs. ETRs based on the combination of KRAB, DNMT3A and DNMT3L effectors domains have been proven to be effective against the large majority of protein-coding genes, with a wide - around 1 kilobase-long - permissive targeting window centered on the transcription start site^32^. However, if, among the gRNAs that you test, none is *per se* able to permanently silence your target gene, you may consider two, non-mutually exclusive, escape strategies: a) switch from transient to stable – e.g., integrating viral vector-based – expression of the ETRs (either the gRNA or the dCas9 fusion constructs or both when adopting the CRISPR/dCas9 technology); b) switch to the alternative ZFP^8^- or TALE^9^-based DNA binding technologies.

### Evaluate the off-target activity of the ETRs

Multiple studies have shown preliminary *in vitro* indications of the specificity of ETRs based on the combination of KRAB, DNMT3A and DNMT3L effectors domains^30,31,32^. However, if among the gRNAs that you test, none of them show a satisfactory specificity profile, in terms of transcriptional regulation and/or de novo DNA methylation, also in this case you may test two non-mutually exclusive strategies: a) reduce the half-life of the ETRs inside the cell. For instance, compared to plasmids, both mRNA and protein delivery are expected to reduce the cell exposure-time to the ETRs and, consequently, the likelihood of off-target activity^55^. Also consider a dose de-escalation approach; b) switch to more recent Cas9 variants optimized to reduce the off-target binding of the platform^56^ or to alternative ZFP^8^- or TALE^9^-based DNA binding technologies. Please consider that compared to mock-treated samples, some of the transcriptional and - less likely - CpG methylation alterations that you can measure in silenced cells may simply derive by the deprivation of your target gene. These should not be considered off-targets of the silencing technology. To identify them, also include in your experimental panel gene disruption by artificial nuclease^8,9,10^ or gene knockdown by RNA interference^33^. Biological alterations due to the functional loss of your target gene should be shared between epigenetic silencing and these alternative technologies.

## AUTHORS CONTRIBUTION

A.M. designed the protocol, conceived and wrote the manuscript with input from all the authors; M.A.C. contributed to design of the protocol and to write the manuscript; S.V., I.M. and D.C. designed the bioinformatics sections of the protocol and revised the manuscript; A.L. designed the protocol, conceived and wrote the manuscript with input from all the authors.

## ACKNOWLDEGMENTS

The authors want to thank Angelo Amabile, Paola Capasso, Ilaria Caserta, Tania Baccega, Alice Reschigna, Valeria Mollica, Alberto Coglot, Deborah Cipria for the collaborative effort at developing the epigenetic silencing technology throughout the years; Dejan Lazarevic and Francesca Giannese for critical review of the RNA-Seq and MeDIP-Seq analyses described in the protocol. Illustrations have been created with BioRender.com. This work was supported by grants to A.L. from Telethon Foundation (TIGET grant no. F1) and the EU Horizon 2020 Program (UPGRADE).

## REFERENCES

1. Cummings et al. Fourteen and counting: unraveling trinucleotide repeat diseases. Hum Mol Genet. 2000 pr 12;9(6):909–16. doi: 10.1093/hmg/9.6.909.

2. Tebas et al. Gene editing of CCR5 in autologous CD4 T cells of persons infected with HIV. N Engl J Med. 2014 Mar 6;370(10):901–10. doi: 10.1056/NEJMoa1300662

3. Frangoul et al. CRISPR-Cas9 Gene Editing for Sickle Cell Disease and β-Thalassemia. N Engl J Med. 2021 Jan 21;384(3):252–260. doi: 10.1056/NEJMoa2031054

4. Murty et al. Gene editing to enhance the efficacy of cancer cell therapies. Mol Ther. 2021 Nov 3;29(11):3153–3162. doi: 10.1016/j.ymthe.2021.10.001.

5. Lanza et al. Engineering universal cells that evade immune detection. Nat Rev Immunol, 2019. Dec;19(12):723–733. doi: 10.1038/s41577-019-0200-1.

6. Matharu et al. Modulating gene regulation to treat genetic disorders. Nat Rev Drug Discov, 2020 Nov;19(11):757–775. doi: 10.1038/s41573-020-0083-7.

7. Sgro et al. Epigenome engineering: new technologies for precision medicine. Nucleic Acids Res. 2020 Dec 16;48(22):12453–12482. doi: 10.1093/nar/gkaa1000.

8. Urnov et al. Genome editing with engineered zinc finger nucleases. Nat Rev Genet. 2010 Sep;11(9):636–46. doi: 10.1038/nrg2842.

9. Joung et al. TALENs: a widely applicable technology for targeted genome editing. Nat. Rev. Mol. Cell Biol. 2013 Jan;14(1):49–55. doi: 10.1038/nrm3486

10. Adli et. al. The CRISPR tool kit for genome editing and beyond. Nat. Commun. 2018 May 15;9(1):1911. doi: 10.1038/s41467-018-04252-2.

11. Gilbert et al. CRISPR-mediated modular RNA-guided regulation of transcription in eukaryotes. Cell. 2013 Jul 18;154(2):442–51. doi: 10.1016/j.cell.2013.06.044.

12. Snowden et al. Gene-specific targeting of H3K9 methylation is sufficient for initiating repression in vivo. Curr. Biol. 2002 Dec 23;12(24):2159–66. doi: 10.1016/s0960-9822(02)01391-x.

13. Chen et al. Construction and validation of the CRISPR/dCas9-EZH2 system for targeted H3K27Me3 modification. Biochem. Biophys. Res. Commun. 2019 Apr 2;511(2):246–252. doi: 10.1016/j.bbrc.2019.02.011.

14. Kwon et al. Locus-specific histone deacetylation using a synthetic CRISPR–Cas9-based HDAC. Nat. Commun. 2017 May 12;8:15315. doi: 10.1038/ncomms15315.

15. Stepper et al. Efficient targeted DNA methylation with chimeric dCas9-Dnmt3a-Dnmt3L methyltransferase. Nucleic Acids Res. 2017 Feb 28;45(4):1703–1713. doi: 10.1093/nar/gkw1112.

16. Ecco et al. KRAB zinc finger proteins. Development, 2017 Aug 1;144(15):2719–2729. doi: 10.1242/dev.132605.

17. Witzgall et al. The Krüppel-associated box-A (KRAB-A) domain of zinc finger proteins mediates transcriptional repression. PNAS. 1994. 1994 May 10;91(10):4514–8. doi: 10.1073/pnas.91.10.4514.

18. Mannini et al. Structure/function of KRAB repression domains: Structural properties of KRAB modules inferred from hydrodynamic, circular dichroism, and FTIR spectroscopic analyses. Proteins, 2006. 2006 Mar 15;62(3):604–16. doi: 10.1002/prot.20792.

19. Friedman et al. KAP-1, a novel corepressor for the highly conserved KRAB repression domain. Genes & Dev. 1996. 10: 2067–2078. doi:10.1101/gad.10.16.2067

20. Iyengar et al. KAP1 Protein: An Enigmatic Master Regulator of the Genome. J Biol Chem, 2011. Jul 29;286(30):26267–76. doi: 10.1074/jbc.R111.252569.

21. Schultz et al. Targeting histone deacetylase complexes via KRAB-zinc finger proteins: the PHD and bromodomains of KAP-1 form a cooperative unit that recruits a novel isoform of the Mi-2α subunit of NuRD. Genes & Dev. 2001. 15: 428–443. doi: 10.1101/gad.869501

22. Schultz et al. SETDB1: a novel KAP-1-associated histone H3, lysine 9-specific methyltransferase that contributes to HP1-mediated silencing of euchromatic genes by KRAB zinc-finger proteins. Genes Dev. 2002 Apr 15;16(8):919–32. doi: 10.1101/gad.973302.

23. Nielsen et al. Interaction with members of the heterochromatin protein 1 (HP1) family and histone deacetylation are differentially involved in transcriptional silencing by members of the TIF1 family. EMBO J. 1999 Nov 15;18(22):6385–95. doi: 10.1093/emboj/18.22.6385.

24. Sripathy et al. The KAP1 Corepressor Functions To Coordinate the Assembly of De Novo HP1-Demarcated Microenvironments of Heterochromatin Required for KRAB Zinc Finger Protein-Mediated Transcriptional Repression. Mol Cell Biol. 2006. 2006 Nov;26(22):8623–38. doi: 10.1128/MCB.00487-06.

25. Jurkowska et al. Structure and function of mammalian DNA methyltransferases.Chembiochem 2011 Jan 24;12(2):206–22.doi: 10.1002/cbic.201000195.

26. Da Jia et al. Structure of Dnmt3a bound to Dnmt3L suggests a model for de novo DNA methylation. Nature. 2007 Sep 13;449(7159):248–51. doi: 10.1038/nature06146

27. Tajima et al. Domain Structure of the Dnmt1, Dnmt3a, and Dnmt3b DNA Methyltransferases. Adv Exp Med Biol. 2016;945:63–86. doi: 10.1007/978-3-319-43624-1_4.

28. Greenberg et al. The diverse roles of DNA methylation in mammalian development and disease. Nat Rev Mol Cell Biol. 2019 Oct;20(10):590–607. doi: 10.1038/s41580-019-0159-6

29. Ishiyama et al. Structure of the Dnmt1 reader module complexed with a unique two-mono-ubiquitin mark on Histone H3 reveals the basis for DNA methylation maintenance. Mol. Cell. 2017. 2017 Oct 19;68(2):350-360.e7. doi: 10.1016/j.molcel.2017.09.037.

30. Amabile et al. Inheritable silencing of endogenous genes by hit-and-run targeted epigenetic editing. Cell. 2016 Sep 22;167(1):219-232.e14. doi: 10.1016/j.cell.2016.09.006

31. Mlambo et al. Designer epigenome modifiers enable robust and sustained gene silencing in clinically relevant human cells. Nucleic Acids Res. 2018 May 18;46(9):4456–4468. doi: 10.1093/nar/gky171.

32. Nuñez et al. Genome-wide programmable transcriptional memory by CRISPR-based epigenome editing. Cell. 2021 Apr 29;184(9):2503-2519.e17. doi: 10.1016/j.cell.2021.03.025.

33. Davidson et al. Current prospects for RNA interference-based therapies. Nat Rev Genet. 2011 May;12(5):329–40. doi: 10.1038/nrg2968.

34. Haapaniemi. CRISPR-Cas9 genome editing induces a p53-mediated DNA damage response. Nat Med. 2018 Jul;24(7):927–930. doi: 10.1038/s41591-018-0049-z.

35. Kosicki et al. Repair of double-strand breaks induced by CRISPR-Cas9 leads to large deletions and complex rearrangements. Nat Biotechnol. 2018 Sep;36(8):765–771. doi: 10.1038/nbt.4192.

36. Ciccia et al. The DNA damage response: making it safe to play with knives. Mol Cell. 2010 2010 Oct 22;40(2):179–204. doi: 10.1016/j.molcel.2010.09.019.

37. Hu et al. Evolved Cas9 variants with broad PAM compatibility and high DNA specificity. Nature. 2018 Apr 5;556(7699):57–63. doi: 10.1038/nature26155.

38. Uhlén M et al. Tissue-based map of the human proteome. Science. 2015. Jan 23;347(6220):1260419. doi: 10.1126/science.1260419.

39. Labun et al. CHOPCHOP v3: expanding the CRISPR web toolbox beyond genome editing. Nucleic Acids Res. 2019 Jul 2;47(W1):W171–W174. doi: 10.1093/nar/gkz365.

40. Kent et al. The human genome browser at UCSC. Genome Res. 2002 Jun;12(6):996–1006. doi: 10.1101/gr.229102.

41. Cheng et al. Multiplexed activation of endogenous genes by CRISPR-on, an RNA-guided transcriptional activator system. Cell Res. 2013 Oct;23(10):1163–71. doi: 10.1038/cr.2013.122.

42. Kabadi et al. Multiplex CRISPR/Cas9-based genome engineering from a single lentiviral vector. Nucleic Acids Res. 2014 Oct 29;42(19):e147. doi: 10.1093/nar/gku749.

43. Dobin A et al. STAR: ultrafast universal RNA-seq aligner. Bioinformatics. 2013 Jan 1;29(1):15–21. doi: 10.1093/bioinformatics/bts635.

44. Liao Y. The Subread aligner: fast, accurate and scalable read mapping by seed-and-vote. Nucleic Acids Res. 2013 May 1;41(10):e108. doi: 10.1093/nar/gkt214.

45. Harrrow J et al. GENCODE: the reference human genome annotation for The ENCODE Project. Genome Res. 2012. Sep;22(9):1760–74. doi: 10.1101/gr.135350.111.

46. Robinson MD et al. A scaling normalization method for differential expression analysis of RNA-seq data. Genome Biol. 2010;11(3):R25. doi: 10.1186/gb-2010-11-3-r25.

47. Robinson MD et al. edgeR: a Bioconductor package for differential expression analysis of digital gene expression data. Bioinformatics. 2010 Jan 1;26(1):139–40. doi: 10.1093/bioinformatics/btp616.

48. Li H et al. Fast and accurate long-read alignment with Burrows-Wheeler transform. Bioinformatics. 2010 Mar 1;26(5):589–95. doi: 10.1093/bioinformatics/btp698.

49. Zhang Y et al. Model-based analysis of ChIP-Seq (MACS). Genome Biol. 2008;9(9):R137. doi: 10.1186/gb-2008-9-9-r137.

50. Quinlan AR et al. BEDTools: The Swiss-Army Tool for Genome Feature Analysis. Curr Protoc Bioinformatics. 2014 Sep 8;47:11.12.1-34. doi: 10.1002/0471250953.bi1112s47.

51. Hansen KD et al. Removing technical variability in RNA-seq data using conditional quantile normalization. Biostatistics. 2012 Apr;13(2):204–16. doi: 10.1093/biostatistics/kxr054

52. Hsu et al. 2013. DNA targeting specificity of RNA-guided Cas9 nucleases. Nat Biotechnol. 2013 Sep;31(9):827–32. doi: 10.1038/nbt.2647.

53. Zeitler et al. Allele-selective transcriptional repression of mutant HTT for the treatment of Huntington’s disease. Nat Med. 2019. 2019 Jul;25(7):1131–1142. doi: 10.1038/s41591-019-0478-3

54. Andersson R et al. Determinants of enhancer and promoter activities of regulatory elements. Nat Rev Genet. 2020 Feb;21(2):71–87. doi: 10.1038/s41576-019-0173-8.

55. Liang X et al. Rapid and highly efficient mammalian cell engineering via Cas9 protein transfection. J Biotechnol. 2015 Aug 20;208:44–53. doi: 10.1016/j.jbiotec.2015.04.024.

56. Han HA et al. Mitigating off-target effects in CRISPR/Cas9-mediated in vivo gene editing. J Mol Med. 2020 May;98(5):615–632. doi: 10.1007/s00109-020-01893-z

