## Supplementary material for "*In vitro* selection of Engineered Transcriptional Repressors for targeted epigenetic silencing and initial evaluation of their specificity profile": Table of Materials

| Item | Vendor | Catalogue Number |
|---|---|---|
| Restriction enzymes | NEB | |
| T4 DNA Ligase | Promega | M1801 |
| Go Taq G2 Hot Start DNA Polymerase | Promega | M7401 |
| One Shot™ TOP10 Chemically Competent cells | ThermoFisher | C404010 |
| NucleoBond Xtra Midi kit for transfection-grade plasmid DNA | Macherey-nagel | 740410.50 |
| K-562 cells | ATCC | CCL-243 |
| Corning™ RPMI 1640 Medium (Mod.) 1X with L-Glutamine | Corning | 10-040-CV |
| CytoFLEX S V4-B4-R3-I2 Flow Cytometer | Beckman Coulter | C01161 |
| BD FACSAria™ Fusion Flow Cytometer | BD Biosciences | |
| 4D-Nucleofector X Unit | Lonza Bioscience | AAF-1003X |
| RNeasy Mini kit | Qiagen | 74106 |
| QIAamp DNA Mini Kit | Qiagen | 51304 |
| 2100 Bioanalizer | Agilent | G2939BA |
| E220 Focused-ultrasonicator | Covari | 500239 |
| MagMeDIP kit | Diagenode | C02010020 |
| IPure kit | Diagenode | C03010011 |
| NextFlex Methylseq kit 1 | Bioo Scientific | 5118-01 |
